# OncoSimulR: genetic simulation of cancer progression with arbitrary epistasis and mutator genes

**DOI:** 10.1101/069500

**Authors:** Ramon Diaz-Uriarte

## Abstract

OncoSimulR implements forward-in-time genetic simulations of diallelic loci in asexual populations with special focus on cancer progression. Fitness can be defined as an arbitrary function of genetic interactions between multiple genes or modules of genes, including epistasis, restrictions in the order of accumulation of mutations, and order effects. Mutation rates can be made to differ between genes, and can be affected by (anti)mutator genes. Also available are sampling from single or multiple simulations, including single-cell sampling, plotting the parent-child relationships of the clones and generating and plotting random fitness landscapes.

**Availability and implementation:** Implemented in R and C++, freely available from BioConductor for Linux, Mac, and Windows under the GNU GPL license. Version 2.3.12 or higher available from: http://www.bioconductor.org/packages/devel/bioc/html/OncoSimulR.html. GitHub repository at: https://github.com/rdiaz02/OncoSimul.

## 1 Introduction

Forward-in-time genetic simulations are widely used in population genetics and cancer research to verify analytic results, to generate data to assess the performance of statistical methods, and to examine complex models that are mathematically intractable (reviews and examples in Diaz-Uriarte, 2015; Thornton, 2014; Yuan *et al.*, 2012). In many scenarios we will want to use a wide range of populations sizes, with possibly large genomes, and with a flexible mechanism to specify the effects of mutations on both fitness and mutation rates (to model mutator/antimutator genes; Gerrish *et al.*, 2007). If the effects of sampling are relevant (e.g Diaz-Uriarte, 2015), we should be able to take samples under different sampling schemes and if understanding the dynamics and history matters, we will also want to track the complete history of the clones. Many forward-in-time simulators are available (see reviews in Peng *et al.*, 2012; Thornton, 2014; Yuan *et al.*, 2012 and the page (https://popmodels.cancercontrol.cancer.gov/gsr/). In the field of evolutionary genetics some of the tools closest to fulfil the above needs are simuPOP (Peng *et al.*, 2012), fwdpp (Thornton, 2014) and FFPopSim (Zanini and Neher, 2012); these programs, however, miss some of the above mentioned features, especially flexible ways to specify fitness and mutator effects, order effects, or gene-specific mutation rates. TTP (Reiter *et al.*, 2013), targeted to cancer progression, is limited to four genes. Motivated by these needs, I have developed OncoSimulR.

## 2 Functionality

### 2.1 Fitness and mutator effects specification

OncoSimulR provides two ways to specify fitness effects:

1. Explicitly mapping genotypes to fitness.
2. Using a lego system to combine:
  a. Effects of individual genes and epistatic effects of any order that involve an arbitrary number of genes.
  b. Order effects involving arbitrary numbers of genes. With order effects, recently discovered in myeloma (Ortmann *et al.*, 2015), the fitness of a genotype with genes A and B mutated differs depending on whether A or B mutated first.
  c. Directed acyclic graphs (DAGs), as used in cancer progression networks such as Oncogenetic Trees and Conjunctive Bayesian Networks (Beerenwinkel *et al.*, 2014), to represent restrictions in the order of accumulation of mutations. How the restriction is enforced and whether conjunctions in the DAG represent AND, OR, or XOR relationships is fully controllable by the user.

We can also define “modules” so that we model the effects on fitness in terms not of individual genes but modules or pathways. Modules can be used with all the lego system specifications (i.e., epistasis, order effects, and DAGs).

Mutator/antimutator genes can be specified similar to fitness effects (though DAGs and order effects are not used to avoid overengineering) and can also involve modules. Genes with (anti)mutator effects can also have direct effects on fitness. Regardless of (anti)mutator genes, mutation rates can be gene-specific or common to all genes.

### 2.2 Running and sampling simulations

We can run simulations using exponential growth (two different types) or a model with carrying capacity that follows McFarland *et al.* (2013). OncoSimulR uses the fast, state-of-the-art, BNB algorithm of Mather *et al.* (2012). Running multiple simulations is parallelized by default in POSIX systems. Simulations need not start from the wildtype genotype: any genotype can be used as the starting point for a simulation. There is no pre-set limit on genome size, and the documentation shows examples with 50000 genes. Simulations can reach total populations sizes > 10^14^.

Simulations can be stopped when total population size, number of time periods, or number of mutated driver genes in any clone are larger than user-specified values. The simulations can also be ended using a stochastic detection mechanism where the probability of detecting a tumor increases with total population size (the parameters controlling this process are modifiable by the user).

Once the simulation is ended, we can take samples from the simulation. It is possible to obtain single-cell samples or whole-tumor samples. In the later case, the “genotype” of the sample will be determined by the frequency at which each gene is mutated in the whole sample (the threshold for detection being set by the user). Regardless of how simulations have been stopped, it is possible to re-sample the history taking samples at uniformly distributed times, or as soon as the population reaches a certain size. As multiple simulations can be run, serially or in parallel, the sampling procedures deal with both single and multiple simulations.

### 2.3 Additional functionality

OncoSimulR provides additional functionality, making it suitable for exploring a wide range of research questions:

- Plotting the parent-child relationships of all clones generated during the simulation, with filtering based on clone size.
- Generating random fitness landscapes using a three-parameter Rough Mount Fuji model that includes the House of Cards and the additive models as special cases (Szendro *et al.*, 2013).
- Plotting fitness landscapes (inspired by MAGELLAN; Brouillet *et al.*, 2015).
- Generating random DAGs that represent restrictions in the order of accumulation of mutations.

### 2.4 Tests and code coverage and documentation

OncoSimulR includes more than 2000 tests that are run routinely at every build/check cycle. These tests provide a code coverage of more than 95% including both the C++ and R code (virtually all non-covered lines correspond to code that should be triggered by as-of-yet-undetected bugs). Another set of 500 long-running (several hours) tests can be run on demand. OncoSimulR includes the standard R function documentation and a comprehensive (more than 140 pages) vignette with fully commented examples of usage.

## 3 Conclusion

Salient features of OncoSimulR compared to other simulators are the unparalleled flexibility to specify fitness and mutator effects and the inclusion of order effects. OncoSimulR can thus be used to address a wide range of questions that span from the effect of mutator/antimutator genes, to the interplay between fitness landscapes, population sizes and mutation rates. OncoSimulR also adds features that are particularly convenient for examining cancer evolution models, such as modules and the flexibility for sampling and stopping the simulations. OncoSimulR is thus of interest for a broad scientific readership that covers from population and evolutionary geneticists to computational oncologists.

## Acknowledgements

C. Lazaro-Perea and A. Parramon for comments on the ms. C. D. McFarland and W. Mather and L. Tsimring for answers about their models/algorithms.

